# Live-cell super-resolution imaging of actin using LifeAct-14 with a PAINT-based approach

**DOI:** 10.1101/2022.11.15.516635

**Authors:** Haresh Bhaskar, Dirk-Jan Kleinjan, Curran Oi, Zoe Gidden, Susan J. Rosser, Mathew H. Horrocks, Lynne Regan

## Abstract

We present direct-LIVE-PAINT, an easy-to-implement approach for the nanoscopic imaging of protein structures in live cells using labeled binding peptides. We demonstrate the feasibility of direct-LIVE-PAINT with an actin-binding peptide fused to EGFP, the location of which can be accurately determined as it transiently binds to actin filaments. We show that direct-LIVE-PAINT can be used to image actin structures below the diffraction-limit of light and have used it to observe the dynamic nature of actin in live cells. We envisage a similar approach could be applied to imaging other proteins within live mammalian cells.

## Introduction

Molecules can be visualized at a resolution below the diffraction-limit of light (~200 nm) using a variety of optical techniques (Hell and Wichmann, 1994, Betzig et al., 2006, Rust et al., 2006) that are collectively grouped under the term super-resolution (SR) microscopy. In single-molecule localization microscopy (SMLM) approaches, the spatiotemporal separation of individual emitters is achieved via the stochastic activation of fluorophores, each of which can be localized with nanometer accuracy (for a comprehensive review of techniques, see (Horrocks et al., 2014). In points accumulation for imaging nanoscale topography (PAINT), fluorescent molecules temporarily bind to surfaces, such as membranes, and are localized allowing for SR imaging (Sharonov and Hochstrasser, 2006). Since the initial demonstration of this strategy, variations on the method have been developed by several different groups (Giannone et al., 2013, Whiten et al., 2018, Sanders et al., 2022). DNA-PAINT has proven particularly useful and has been applied to imaging DNA origami arrays *in vitro*, and proteins in fixed samples (Schnitzbauer et al., 2017, Guo et al., 2019). As DNA-PAINT relies on tagging biomolecules with unique oligonucleotide sequences, it can also be used for multiplexed imaging (Jungmann et al., 2014). An analogous approach, peptide-PAINT, has also been developed using peptide-peptide interaction pairs (Eklund et al., 2020).

The requirement for fixation and permeabilization, however, generally precludes the use of either DNA-PAINT or peptide-PAINT for live-cell imaging, although in some cases DNA-PAINT has been used to image membrane proteins on the outer surface of live cells (Strauss et al., 2018). To circumvent the limitation of cell fixation and permeabilization, we recently developed LIVE-PAINT (Live cell Imaging using reVersible intEractions-PAINT), a method that uses transient protein-protein interactions to perform SR imaging in live cells (Oi et al., 2020). In this approach, the protein-of-interest is genetically fused to a short peptide sequence, and a SR image is generated as this is transiently bound by an interacting peptide fused to a fluorescent protein (Oi et al., 2020). While useful for imaging targets in live cells, the LIVE-PAINT technique has only been demonstrated in live yeast, and also requires the modification of the target protein, which may perturb its structure and function. To circumvent this limitation, we introduce direct-LIVE-PAINT. Rather than relying on tagging the protein-of-interest with a peptide sequence, direct-LIVE-PAINT uses genetically-encoded fluorescent peptides that directly interact with the unmodified, endogenous protein, enabling SR imaging in live mammalian cells.

Actin is an abundant protein that is a component of the eukaryotic cytoskeleton. Its dynamic conversion between the monomeric (G-actin) and the filamentous (F-actin) state underlies key cellular functions. New ways to image F-actin in live cells are especially useful, since when directly fused to a fluorescent protein, actin is not fully functional. One probe, termed LifeAct, uses the N-terminal 17 amino acids of an actin-binding protein Abp140 fused to EGFP to label actin. Although widely used to label actin in live and fixed cells, artifacts have been noted with LifeAct (Kiuchi et al., 2015, Flores et al., 2019). Most notably, it has been reported that LifeAct exhibits a higher affinity for G-actin than for F-actin and that this differential affinity perturbs the equilibrium, causing artifacts (Courtemanche et al., 2016, Flores et al., 2019). Kumari et al. recently reported that a peptide lacking the C-terminal 3 amino acids of LifeAct, which they named LifeAct-14, binds to F-actin with similar affinity (K_D_=1.2 μM) to LifeAct, and does not exhibit preferential binding to G-actin (Kumari et al., 2020). They therefore reasoned it would be less perturbing to cellular function (actin assembly) than LifeAct itself (Figure 1A).

**Figure 1:**
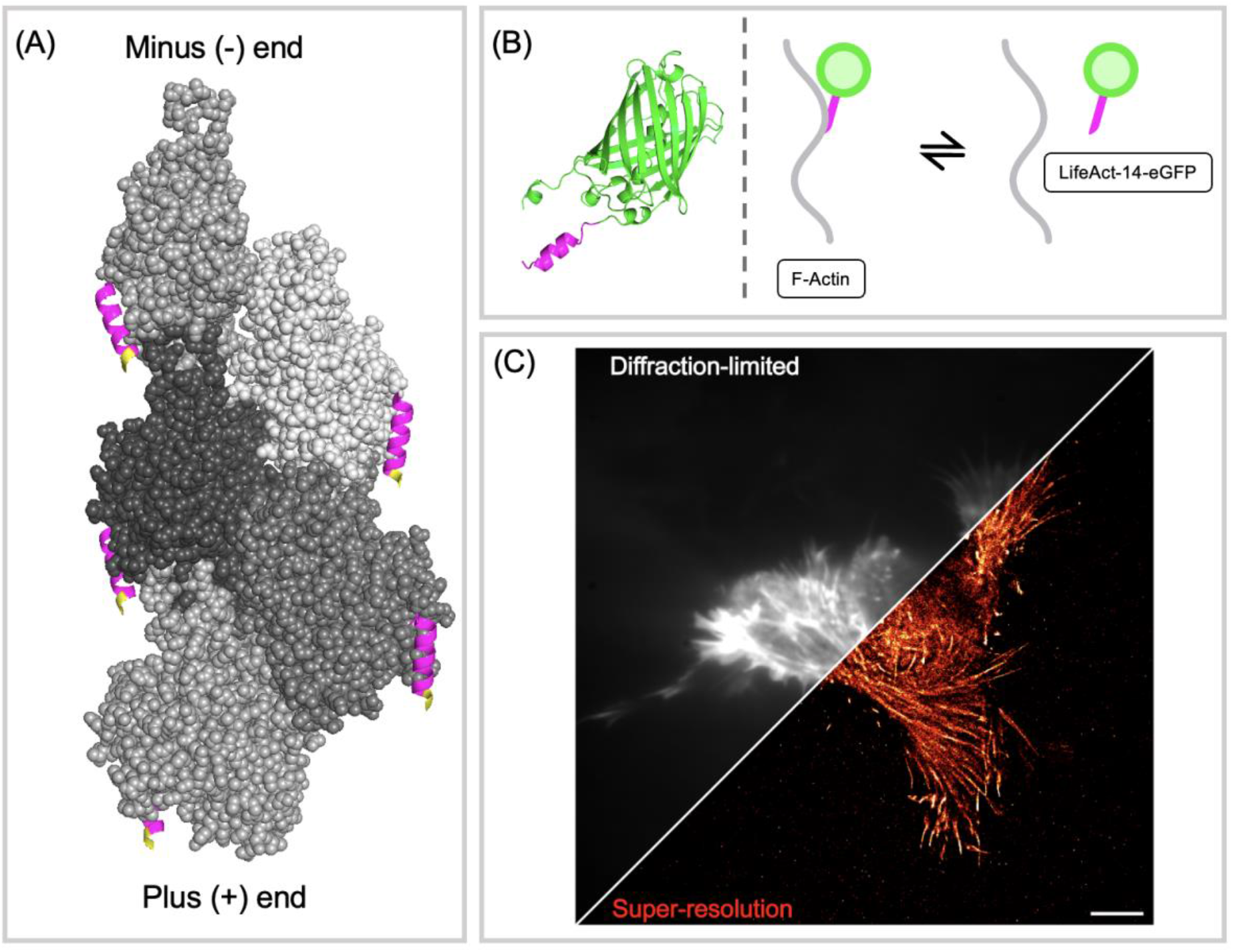
(A) Cryo-EM structure of LifeAct (17 AA) bound to F-actin. Five monomers of actin are shown in space filling representation in shades of gray. Each actin monomer is shown bound to the LifeAct peptide, which is shown in magenta in ribbon representation. The three residues at the C-terminus, colored yellow, indicate the amino acids that are missing from LifeAct-14. Structure 7AD9 retrieved from the PDB (Belyy et al., 2020). (B) Cartoons showing a predicted structure of the LifeAct-14-EGFP construct (LifeAct-14, magenta; EGFP, green) (Jumper et al., 2021) and illustrating the reversible nature of the binding between LifeAct-14-EGFP and actin filaments. EGFP sequence retrieved from FPbase (Lambert, 2019). Structures visualized using PyMOL (The PyMOL Molecular Graphics System, Version 2.0 Schrödinger, LLC.). (C) A HEK293 cell, transiently transfected with a plasmid expressing LifeAct-14-EGFP. The distinctive actin filament structure is clearly visible. The diagonal line separates diffraction limited and super-resolution (SR) imaging. SR image colored ‘red hot’. Diffraction-limited image was obtained by Z-projecting the time-course images (100s, 50ms exposure). Scale bar is 5 μm.

Based on its low micromolar affinity and negligible perturbative effects, we postulated that expression of LifeAct-14-EGFP fusion could be used to obtain SR images of filamentous actin via the direct-LIVE-PAINT approach (Figure 1B,C). As with LIVE-PAINT, we show that direct-LIVE-PAINT is dependent on the concentration of the labeled peptide and determine the optimum expression conditions for LifeAct-14-EGFP in HEK-293 cells. We also demonstrate that it is possible to image the dynamic actin structure in live cells, achieving a spatial resolution of 80 nm, with a temporal resolution of 12.5 s. By enabling the tracking of actin dynamics in live cells, we believe that direct-LIVE-PAINT with LifeAct-14-EGFP will serve as a valuable tool to improve our current understanding of the role played by the cytoskeleton in motility, adhesion and other cellular processes. Furthermore, direct-LIVE-PAINT presents a method for probing a range of biomolecules in SR, provided the peptide has suitable binding kinetics for the target-of-interest.

## Results

### Transfection of LifeAct-14-EGFP allows the super-resolution imaging of actin

As with any SMLM-based technique, the success of direct-LIVE-PAINT relies on having the optimum levels of active fluorophore to enable spatiotemporal separation. We therefore investigated how varying the transfection levels of LifeAct-14-EGFP affected our ability to distinguish individual fluorophores (Figure 2). At low transfection levels (0.032-0.8 ng DNA), negligible binding was observed. At higher concentrations (4 ng DNA), however, we observed transient binding of LifeAct-14-EGFP, allowing the localization of individual emitters (54k locs). At higher concentrations still (20 ng DNA), the binding rate increased, leading to higher density SR images (186k localizations) and clear filament structure (spatial resolution of 100 nm as determined by Fourier Ring Correlation (FRC)) (ten Brink, T. RustFRC [Computer software]). At higher expression levels (100 ng DNA) however, individual binding events could no longer be observed due to saturation of the actin-binding sites with LifeAct-14-EGFP. Based on these observations, a transfection concentration of 20 ng was optimum for LifeAct-14-EGFP in live HEK-293 cells. The images at each transfection amount (Figure 2) are representative examples seen in ~60-70% of the transfected cell population (data not shown).

**Figure 2:**
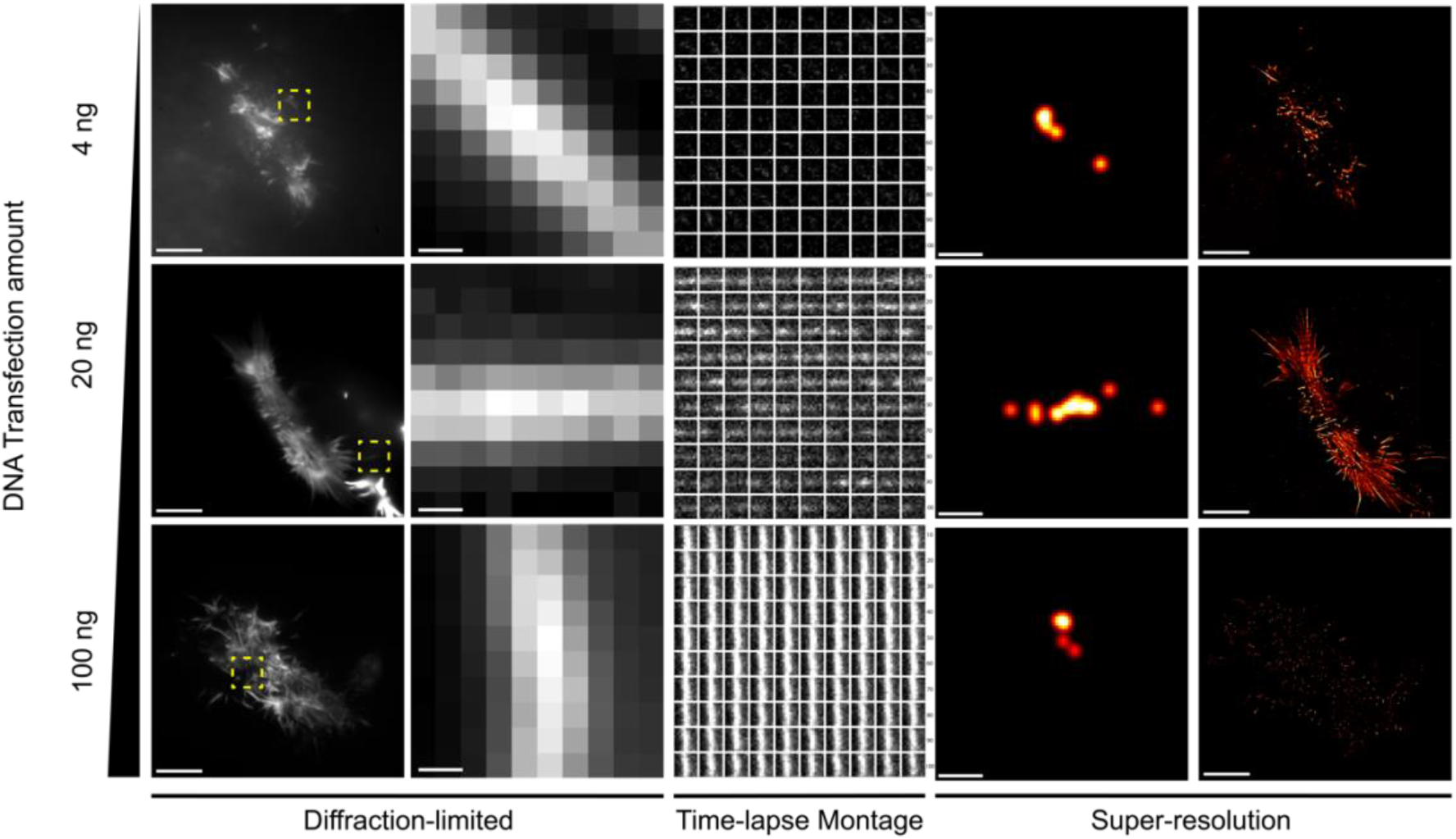
Intensity fluctuations and localizations of LifeAct-14-EGFP targeting F-actin in HEK293 cells in response to a range of DNA transfection amounts. Diffraction-limited images shown on the left are median intensity Z-projections where the first column shows an entire field of view (FOV) taken over 2000 frames (100 s, 50 ms exposure, 10 μm scale bar) and the adjacent column shows a region of interest (ROI from the region in the dashed box shown in yellow over a 100-frame subset. The time-lapse montage column shows intensity fluctuations within the ROI across the 100-frame subset (time - left to right and top to bottom). The SR images constructed from the detected localizations in the ROI (0.2 μm scale bar) and then across the entire FOV (10 μm scale bar, 2000 frames) are shown in the far right. Precision threshold < 30 nm.

### Actin filaments in live cells are dynamic structures

Actin is a key component of the cytoskeleton and is responsible for maintaining cell structure and shape. Due to the non-perturbative nature of direct-LIVE-PAINT, and its ability to image in live cells, we used it to track the dynamics of actin filaments.

The spatial resolution of images generated using SMLM is dependent on both the precision of each emitter localized, and the density of the localizations. To achieve the highest resolution, it can therefore be necessary to image over several minutes, resulting in the loss of temporal resolution. Thus, there is a compromise to be made between achieving the highest spatial and temporal resolution. We therefore investigated how the imaging time affected the spatial resolution of direct-LIVE-PAINT by analyzing subsets of frames generated from LifeAct-14-EGFP binding to actin in live HEK-293 cells (Figure 3). We set a more stringent precision threshold (< 20 nm) to determine whether spatial resolution improves, while still detecting sufficient localizations to discern filament structure. With increasing imaging time (example images in Figure 3A), the spatial resolution increased from ~100 nm to 80 nm (Figure 3B), and more detail was evident in the images. As localizations were accumulated over time (Figure 3B), clearer structure was observed. At higher integration times, the resolution appeared to plateau. To increase the number of localizations further, we applied a less stringent precision threshold (< 30 nm). Although this reduces the spatial resolution (80 nm to 100 nm), the increase in the number of localizations (60k vs 186k) allowed the movement of filaments to be visualized at a time-resolution of 12.5 s. Indeed, this can be observed in the time-montage of the direct-LIVE-PAINT images in a ROI (Figure 3C), where the filament (dashed white box) clearly changes in shape over 50 s. The SR plot in which the localizations were colored according to the time they were detected (Figure 3D) further illustrates this behavior; the dynamics of the same filament (red box) can be observed throughout the imaging time-course.

**Figure 3:**
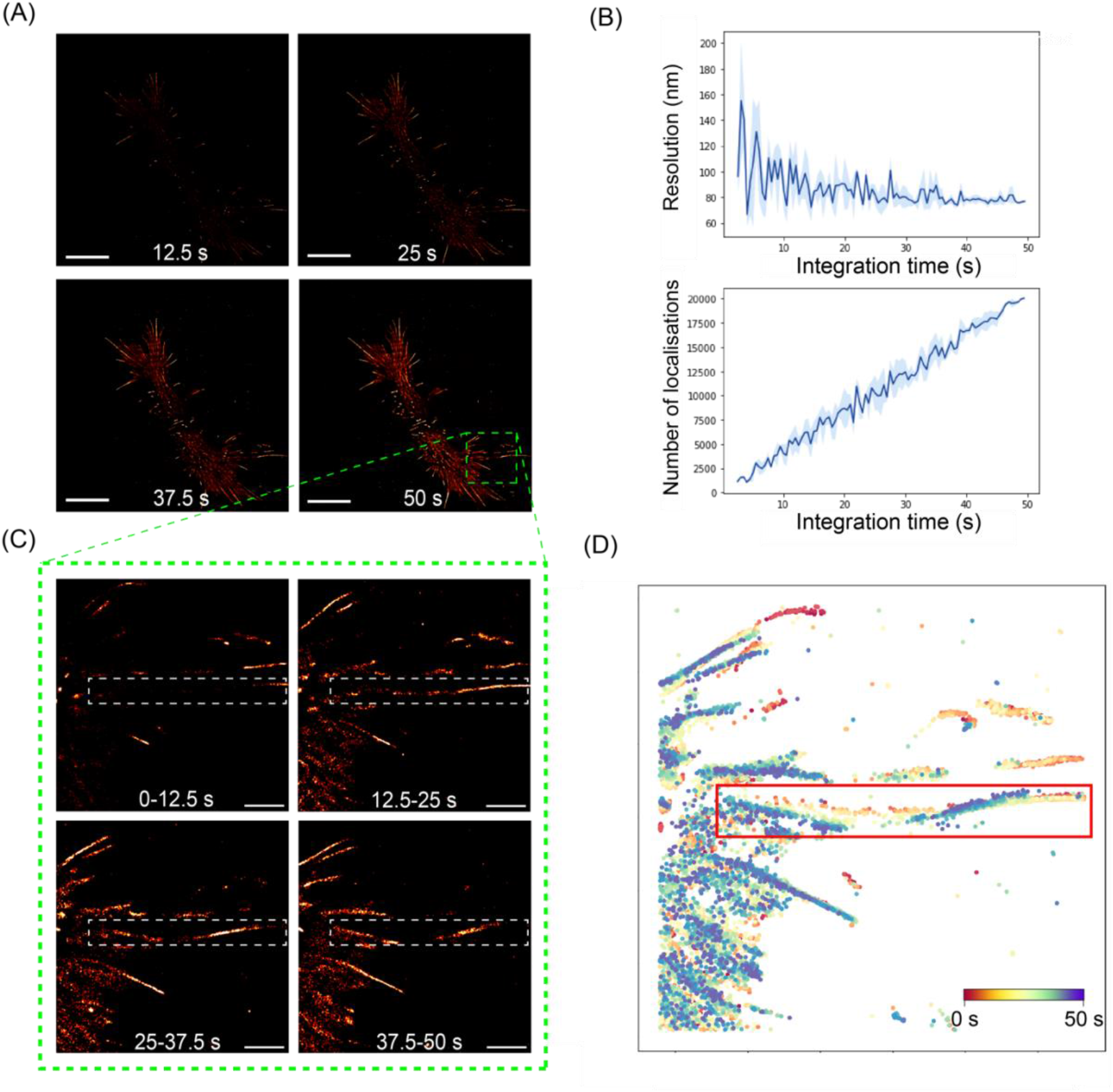
F-actin structures are dynamic within timescales of imaging and can be detected through direct-LIVE-PAINT. A) SR images of F-actin imaged with LifeAct-14-EGFP at a range of integration times in a HEK293 cell. 10 μm scale bar. (B) Resolution and localization number in response to increasing integration times. Frame subsets were sampled randomly at each integration time for resolution and localisation number calculation. Standard deviation error shown as shaded region (n=3). Precision threshold < 20 nm. (C) SR images shown as a time montage in a ROI (dashed green box from (A)), tracking an actin filament (dashed white box) through the same timeframe. 2 μm scale bar. (D) Map of localizations color-coded based on time of acquisition across the same timeframe. 2.5 s split, time scales from red, to yellow, to green, to blue. Red box indicates the same filament-of-interest that changes in shape during this timeframe.

## Conclusion

In summary, we demonstrate that direct-LIVE-PAINT can be used to probe protein targets in live cells. We used direct-LIVE-PAINT with LifeAct14-EGFP to characterize the structure and dynamics of F-actin at the nanoscale. This is the first LIVE-PAINT-based imaging in mammalian cells. Although shown with an actin-binding peptide here, other classes of peptides such as nanobodies (Muyldermans et al., 2009), affibodies (Nord et al., 1997), and DARPins (Binz et al., 2003) that target various proteins in mammalian cells could potentially be used with direct-LIVE-PAINT.

## Materials and Methods

### DNA amplification and purification

LifeAct-14-EGFP plasmid was received as agar stabs (#158750, Addgene) (Kumari et al., 2020). Cultures were grown up according to standard protocols. Plasmid DNA was purified using a QIAGEN Spin Miniprep Kit according to manufacturer protocols and concentration and purity was assessed on a NanoDrop™ Microvolume spectrophotometer. Extracted DNA was stored at −20°C until ready for transfection.

### Mammalian cell culture and transfection

HEK293 cells were cultured according to standard protocols (ATCC). Briefly, cells were cultured in DMEM supplemented with 10% FBS, 1% L-glutamine and 1% Pen/Strep in T25 flasks. Once ~70% confluent, cells were detached with TrypLE (Gibco) by adding 0.5 mL and incubating for 2-3 mins in the incubator at 37°C. Then, 4 mL of low-fluorescence DMEM, without phenol Red (Gibco™, Cat no. 21063-029) was added and the cell density of the suspension was counted with an automated cell counter (Countess II, Thermofisher Scientific) using 0.4% Trypan Blue at a 1:1 cell:dye ratio (1:2 dilution) Ibidi imaging plates (#81817) were seeded at 15k cells in 100 *μ*L per well (1.5e5 cells/ mL) and allowed to adhere for ~20 mins at room temperature.

Meanwhile transfection reagents (Lipofectamine ™ 3000 Transfection Reagent, Invitrogen™) were made up according to the manufacturer’s protocol. Briefly, the P3000 reagent was diluted into Opti-MEM™ (Gibco™) and mixed with different amounts of DNA based on the transfection condition. Next, Lipofectamine 3000 Reagent diluted in Opti-MEM was added and mixed thoroughly by pipeting up and down. The lipofectamine-DNA mixture was incubated for 10 mins at room temperature before being added to the cells at 25 *μ*L per well. Plates were incubated for 24h at 37° C, under 5% CO2, prior to imaging.

Because this is a proof of principle experiment, we were satisfied with empirically optimizing the DNA amount used in transfection to identify the amount that gave appropriate binding behavior to allow SMLM.

### TIRF Microscopy

All imaging was performed based on a previously published protocol (Oi et al., 2020). Briefly, a home built TIRF instrument using an inverted Nikon^®^ TI2 microscope was used with a heated stage incubator to maintain humidity and temperature at 37°C. EGFP was excited with a laser light at a wavelength of 488 nm and the emission was collected by the same objective by separating it from the returning TIR beam with the use of a dichroic mirror. Images were acquired at an exposure of 50 ms and at a laser power density of 3.5 W/cm^2^.

Fewer localizations were expected from initial frames as the sample bleached down to a suitable labeling density. ImageJ was used for all data acquisition and analysis along with the Micromanager software for microscope automation.

All SR image construction was performed using the FIJI (ImageJ2; Version 2.3.0) plug-in ThunderSTORM (version dev-2016–09-10-b1) by running the *Run analysis* command. Localizations were visualized using the *Normalized Gaussian* method (width=precision) along with a < 30 nm precision threshold, unless otherwise stated.

Resolution and localization number calculations were performed using the RustFRC python package (ten Brink, T. RustFRC [Computer software]) and the script is available at https://doi.org/10.5281/zenodo.7290477. Briefly, consecutive frame subsets of a specific size, determined by the integration time, were randomly sampled from the total frames dataset for resolution and localisation number calculation.

## Author Contributions

HB, MHH and LR led the study, designed the experiments, and wrote the manuscript. HB carried out all the experiments with guidance from DJK & SR in cell culture and MHH in imaging and analysis. CO and ZG offered guidance in experiments and analysis. All authors provided feedback on the manuscript.

## Acknowledgements

HB acknowledges the support of Wellcome Trust via the Integrative Cell Mechanisms PhD program (grant number 226437/Z/22/Z). The super-resolution instrument was funded by UCB Biopharma and a kind donation from Dr. Jim Love. We offer special thanks to members of the Rosser Lab for their tissue culture expertise, Dr Rebecca Saleeb for help with the SMLM analysis workflow and Dr Raef Shams for help with protein structure visualization.

## References

Belyy, A., Merino, F., Sitsel, O. & Raunser, S. (2020) Structure of the Lifeact–F-actin complex C.A. Parent (ed.). PLOS Biology. 18 (11), e3000925. doi:10.1371/journal.pbio.3000925.

Betzig, E., Patterson, G.H., Sougrat, R., Lindwasser, O.W., Olenych, S., Bonifacino, J.S., Davidson, M.W., Lippincott-Schwartz, J. & Hess, H.F. (2006) Imaging Intracellular Fluorescent Proteins at Nanometer Resolution. Science. 313 (5793), 1642–1645. doi:10.1126/science.1127344.

Binz, H.K., Stumpp, M.T., Forrer, P., Amstutz, P. & Plückthun, A. (2003) Designing Repeat Proteins: Well-expressed, Soluble and Stable Proteins from Combinatorial Libraries of Consensus Ankyrin Repeat Proteins. Journal of Molecular Biology. 332 (2), 489–503. doi:10.1016/S0022-2836(03)00896-9.

Clowsley, A.H., Kaufhold, W.T., Lutz, T., Meletiou, A., Di Michele, L. & Soeller, C. (2021) Repeat DNA-PAINT suppresses background and non-specific signals in optical nanoscopy. Nature Communications. 12 (1), 501. doi:10.1038/s41467-020-20686-z.

Courtemanche, N., Pollard, T.D. & Chen, Q. (2016) Avoiding artefacts when counting polymerized actin in live cells with LifeAct fused to fluorescent proteins. Nature Cell Biology. 18 (6), 676–683. doi:10.1038/ncb3351.

Eklund, A.S., Ganji, M., Gavins, G., Seitz, O. & Jungmann, R. (2020) Peptide-PAINT Super-Resolution Imaging Using Transient Coiled Coil Interactions. Nano Letters. 20 (9), 6732–6737. doi:10.1021/acs.nanolett.0c02620.

Flores, L.R., Keeling, M.C., Zhang, X., Sliogeryte, K. & Gavara, N. (2019) Lifeact-TagGFP2 alters F-actin organization, cellular morphology and biophysical behaviour. Scientific Reports. 9 (1), 3241. doi:10.1038/s41598-019-40092-w.

Giannone, G., Hosy, E., Sibarita, J.-B., Choquet, D. & Cognet, L. (2013) High-Content Super-Resolution Imaging of Live Cell by uPAINT. In: A.A. Sousa & M.J. Kruhlak (eds.). Nanoimaging. Methods in Molecular Biology. Totowa, NJ, Humana Press. pp. 95–110. doi:10.1007/978-1-62703-137-0_7.

Guo, S.-M., Veneziano, R., Gordonov, S., Li, L., Danielson, E., Perez de Arce, K., Park, D., Kulesa, A.B., Wamhoff, E.-C., Blainey, P.C., Boyden, E.S., Cottrell, J.R. & Bathe, M. (2019) Multiplexed and high-throughput neuronal fluorescence imaging with diffusible probes. Nature Communications. 10 (1), 4377. doi:10.1038/s41467-019-12372-6.

Hell, S.W. & Wichmann, J. (1994) Breaking the diffraction resolution limit by stimulated emission: stimulated-emission-depletion fluorescence microscopy. Optics Letters. 19 (11), 780. doi:10.1364/OL.19.000780.

Horrocks, M.H., Palayret, M., Klenerman, D. & Lee, S.F. (2014) The changing point-spread function: single-molecule-based super-resolution imaging. Histochemistry and Cell Biology. 141 (6), 577–585. doi:10.1007/s00418-014-1186-1.

Jumper, J., Evans, R., Pritzel, A., Green, T., Figurnov, M., et al. (2021) Highly accurate protein structure prediction with AlphaFold. Nature. 596 (7873), 583–589. doi:10.1038/s41586-021-03819-2.

Jungmann, R., Avendaño, M.S., Woehrstein, J.B., Dai, M., Shih, W.M. & Yin, P. (2014) Multiplexed 3D cellular super-resolution imaging with DNA-PAINT and Exchange-PAINT. Nature Methods. 11 (3), 313–318. doi:10.1038/nmeth.2835.

Kiuchi, T., Higuchi, M., Takamura, A., Maruoka, M. & Watanabe, N. (2015) Multitarget super-resolution microscopy with high-density labeling by exchangeable probes. Nature Methods. 12 (8), 743–746. doi:10.1038/nmeth.3466.

Kumari, A., Kesarwani, S., Javoor, M.G., Vinothkumar, K.R. & Sirajuddin, M. (2020) Structural insights into actin filament recognition by commonly used cellular actin markers. The EMBO Journal. 39 (14). doi:10.15252/embj.2019104006.

Lambert, T.J. (2019) FPbase: a community-editable fluorescent protein database. Nature Methods. 16 (4), 277–278. doi:10.1038/s41592-019-0352-8.

Lelek, M., Gyparaki, M.T., Beliu, G., Schueder, F., Griffié, J., Manley, S., Jungmann, R., Sauer, M., Lakadamyali, M. & Zimmer, C. (2021) Single-molecule localization microscopy. Nature Reviews Methods Primers. 1 (1), 39. doi:10.1038/s43586-021-00038-x.

Muyldermans, S., Baral, T.N., Retamozzo, V.C., De Baetselier, P., De Genst, E., Kinne, J., Leonhardt, H., Magez, S., Nguyen, V.K., Revets, H., Rothbauer, U., Stijlemans, B., Tillib, S., Wernery, U., Wyns, L., Hassanzadeh-Ghassabeh, Gh. & Saerens, D. (2009) Camelid immunoglobulins and nanobody technology. Veterinary Immunology and Immunopathology. 128 (1–3), 178–183. doi:10.1016/j.vetimm.2008.10.299.

Nord, K., Gunneriusson, E., Ringdahl, J., Ståhl, S., Uhlén, M. & Nygren, P.-Å. (1997) Binding proteins selected from combinatorial libraries of an α-helical bacterial receptor domain. Nature Biotechnology. 15 (8), 772–777. doi:10.1038/nbt0897-772.

Oi, C., Gidden, Z., Holyoake, L., Kantelberg, O., Mochrie, S., Horrocks, M.H. & Regan, L. (2020a) LIVE-PAINT allows super-resolution microscopy inside living cells using reversible peptide-protein interactions. Communications Biology. 3 (1), 458. doi:10.1038/s42003-020-01188-6.

Oi, C., Mochrie, S.G.J., Horrocks, M.H. & Regan, L. (2020b) PAINT using proteins: A new brush for super-resolution artists. Protein Science. 29 (11), 2142–2149. doi:10.1002/pro.3953.

Riedl, J., Crevenna, A.H., Kessenbrock, K., Yu, J.H., Neukirchen, D., Bista, M., Bradke, F., Jenne, D., Holak, T.A., Werb, Z., Sixt, M. & Wedlich-Soldner, R. (2008) Lifeact: a versatile marker to visualize F-actin. Nature Methods. 5 (7), 605–607. doi:10.1038/nmeth.1220.

Rust, M.J., Bates, M. & Zhuang, X. (2006) Sub-diffraction-limit imaging by stochastic optical reconstruction microscopy (STORM). Nature Methods. 3 (10), 793–796. doi:10.1038/nmeth929.

Sanders, E.W., Carr, A.R., Bruggeman, E., Körbel, M., Benaissa, S.I., Donat, R.F., Santos, A.M., McColl, J., O’Holleran, K., Klenerman, D., Davis, S.J., Lee, S.F. & Ponjavic, A. (2022) resPAINT: Accelerating Volumetric Super-Resolution Localisation Microscopy by Active Control of Probe Emission**. Angewandte Chemie International Edition. 61 (42). doi:10.1002/anie.202206919.

Schnittler, H., Taha, M., Schnittler, M.O., Taha, A.A., Lindemann, N. & Seebach, J. (2014) Actin filament dynamics and endothelial cell junctions: the Ying and Yang between stabilization and motion. Cell and Tissue Research. 355 (3), 529–543. doi:10.1007/s00441-014-1856-2.

Schnitzbauer, J., Strauss, M.T., Schlichthaerle, T., Schueder, F. & Jungmann, R. (2017) Super-resolution microscopy with DNA-PAINT. Nature Protocols. 12 (6), 1198–1228. doi:10.1038/nprot.2017.024.

Sharonov, A. & Hochstrasser, R.M. (2006) Wide-field subdiffraction imaging by accumulated binding of diffusing probes. Proceedings of the National Academy of Sciences. 103 (50), 18911–18916. doi:10.1073/pnas.0609643104.

Strauss, S., Nickels, P.C., Strauss, M.T., Jimenez Sabinina, V., Ellenberg, J., Carter, J.D., Gupta, S., Janjic, N. & Jungmann, R. (2018) Modified aptamers enable quantitative sub-10-nm cellular DNA-PAINT imaging. Nature Methods. 15 (9), 685–688. doi:10.1038/s41592-018-0105-0.

Whiten, D.R., Zuo, Y., Calo, L., Choi, M.-L., De, S., Flagmeier, P., Wirthensohn, D.C., Kundel, F., Ranasinghe, R.T., Sanchez, S.E., Athauda, D., Lee, S.F., Dobson, C.M., Gandhi, S., Spillantini, M.-G., Klenerman, D. & Horrocks, M.H. (2018) Nanoscopic Characterisation of Individual Endogenous Protein Aggregates in Human Neuronal Cells. ChemBioChem. 19 (19), 2033–2038. doi:10.1002/cbic.201800209.

